# RNA-sequencing Strain-specific Genome Alignment Increases Differential Expression Findings in Comparison of C57BL/6J and DBA/2J Nucleus Accumbens

**DOI:** 10.1101/2025.05.25.656014

**Authors:** ER Gnatowski, D Zellif, MG Dozmorov, MF Miles

## Abstract

Alcohol Use Disorder (AUD) is a polygenic disease defined by the inability to regulate alcohol consumption despite adverse consequences. C57BL/6J (B6) and DBA/2J (D2) mice, the progenitor strains to the BXD recombinant inbred strain, exhibit differences in voluntary ethanol consumption and other ethanol behaviors, making them frequently used models for studying genetic influences on ethanol responses. The B6 genome is the standard reference genome for the majority of mouse RNA-sequencing (RNAseq) studies, including studies on D2 mice. We hypothesized that aligning B6 and D2 RNAseq data to their strain specific genome would allow improved detection of differentially expressed genes (DEGs) in comparison of brain gene expression between these two strains. RNA samples obtained from B6 and D2 nucleus accumbens (NAc) tissue were analyzed using a standard RNAseq analysis pipeline except from genome alignment. Following quality control, samples were aligned to either the B6 reference genome (Release 108) or the D2 samples were aligned to a recent homologous genome assembly (GCA_921998315.2). Alignment of D2 samples to the D2 genome showed significantly higher alignment compared to the B6 reference genome (93.82% vs 92.02%, p = 0.0272), but also showed a decrease in the number of total reads assigned (72.30%% vs 74.36%, p <0.0001). When comparing B6 and D2 expression, using the D2 alignment resulted in large increases in the number of differentially expressed genes (DEGs) (10,777 vs 6,191) and differentially utilized exons (DUEs) (81,206 vs 21,223) with resulting changes in gene ontology functional analyses. The gene ontology identified substantial overlap between the two analyses while also adding novel categories. These studies highlight the importance of using strain-specific alignment in increasing the number of reads aligned and the number of DEGs and DUEs identified. The use of strain-specific alignment in RNA-seq studies may provide greater accuracy in investigating gene expression and pathways regulated by ethanol in model organism studies on molecular mechanisms of AUD.

## Introduction

RNA-sequencing (RNAseq) is a widely used technique that quantifies gene expression, serving as a crucial link between the genome and the phenome. This technique involves sequencing RNA fragments and then computationally mapping these sequences back to their original genomic locations, a process known as read alignment. The accuracy of read alignment is critical as it directly impacts the subsequent analysis and interpretation of the data such as gene and exon-level expression quantification, variant calling, and alternative splicing analyses [1; 2; 3; 4]. This accuracy is dependent on several factors including: the choice of reference genome, the precision of alignments as dictated by the quality of the aligner, and the ability to account for sequencing errors and repeated sequences within the genome. Inaccuracies that occur as a result of improper characterization of sample reads ultimately spread to downstream analyses such as the detection of differentially expressed genes, exon utilization and splicing analysis, quantitative trait loci (QTL), gene ontology or functional enrichment [5], and expression network analyses.

Recombinant inbred strains have been widely used in studying mouse models of alcohol use disorder (AUD), as they provide a defined genetic background that facilitates identification of genetic factors influencing alcohol-related behaviors. The BXD recombinant inbred panel was generated by crossbreeding C57BL/6J (B6) and DBA/2J (D2) inbred strains [6]. BXD mice provide an excellent framework for studying alcohol-related phenotypes given the distinct responses to acute and chronic ethanol exposure in B6 versus D2 mice [7; 8; 9]. B6 and D2 mice are genetically very distinct, with differences at approximately 5 million sites in their genomes. This genetic diversity makes them an invaluable resource for analyzing complex phenotypes [10]. The Miles laboratory previously used BXD mice to perform behavioral and expression genetic studies on the acute anxiolytic-like response to ethanol [11]. That work identified *Etanq1*, a robust behavioral QTL on Chromosome 12 for ethanol anxiolytic-like activity and led to defining Ninein as a candidate gene for that QTL. Given the extensive behavioral, physiological, and anatomical phenotyping of these strains as well as their recombinant inbred offspring, it is important that RNA-seq studies examine gene expression differences within these strains most accurately reflect biological reality. The B6 mice were the DNA source for the first high-quality draft sequence of the mouse genome, making their genome one of the most extensively studied and commonly used as the standard alignment sequence for genomic analyses [12]. Thus, the B6 genome has predominantly served as the reference genome for a substantial number of mouse studies, including virtually all RNA-Seq analyses regardless of strain, due to its superior quality and comprehensive annotation.

We therefore hypothesized that employing strain-specific alignment could yield more accurate results that truly reflect the biological differences between the C57BL/6J and DBA/2J mouse strains. In this study, we used basal mRNA from nucleus accumbens tissue for RNA-seq analysis. The current study demonstrates that parental strain alignment increases the number of significant differentially expressed genes and subsequently, changes the biological functions identified by functional enrichment analysis providing new insight into functional differences between B6 and D2 mice.

## Materials and Methods

### Animal Subjects

Male C57BL/6J and DBA/2J mice (n = 5/strain) were obtained from the Jackson Laboratory (Bar Harbor, ME, USA) at 7-8 weeks of age and habituated to the vivarium for at least 1 week prior to initiating experimental studies. Animals were group-housed (four animals per group with ad libitum access to food and water, under a 12-hour light/dark cycle, in a 21 °C environment). Experiments were done in the light phase. All experiments were approved by the Institutional Animal Care and Use Committee of Virginia Commonwealth University and followed the National Institutes of Health Guidelines for the Care and Use of Laboratory Animals. At 9 weeks of age, animals were sacrificed by cervical dislocation. Immediately thereafter, brains were extracted and chilled for 1 minute in cold phosphate buffered saline before brain regions being micro-dissected as previously described (Kerns et al., 2005). Excised regions were placed in individual tubes, flash-frozen in liquid nitrogen, and stored at -80 C. Nucleus Accumbens (NAc), sections were used for RNA extraction and analysis as below.

### RNA extractions

Samples were homogenized with a Polytron® (Kinematica AG, Malters, Switzerland) and total RNA was extracted using a guanidine/phenol/chloroform method (STAT-60, Tel-Test, Inc. Friendswood, TX, United States) as per manufacturer guidelines. Each RNA liquid layer was added to a miRNeasy Mini Column (Cat #: 217004, Qiagen, Hilden, Germany) for cleanup and elution of total RNA. RNA concentration was determined by measuring absorbance at 260 nm using a Nanodrop 2000 (Thermo Scientific). RNA quality and purity was assessed by 260/280 absorbance ratios and by electrophoresis using the Agilent 2100 Bioanalyzer (Agilent Technologies, Savage, MD, United States). Samples all had RNA Integrity Numbers (RIN) ≥ 8 and 260/280 ratios between 1.96 and 2.05.

### RNAseq Library Preparation and Sequencing

RNAseq data have been deposited with the Gene Expression Omnibus resource (GEO accession #GSE274854). Library preparation and sequencing were completed by the VCU Genomics Core. A total of 5 biological replicates were obtained from each strain. Preparation of cDNA libraries was conducted using the standard protocols for the Illumina Stranded mRNA Prep, Ligation Kit (#20040534, Illumina, San Diego, CA, United States). Library insert size was determined using an Agilent Bioanalyzer. Libraries were sequenced on the Illumina NextSeq2000 with 150 bp paired-end reads for a target depth of 100 million reads per sample. A summary of RNAseq metrics can be found in **Table 1**.

**Table 1.** RNA-sequencing information. . RNA quality control metrics, alignment percentages to the reference genome, and count generation analytics.

Figures/Tables
| Sample | RIN Value | Uniquely Mapped Alignment % | Assigned % |
| --- | --- | --- | --- |
| B14 N | 9 | 92.90% | 74.40% |
| B21 N | 9.1 | 91.60% | 74.90% |
| B24 N | 9.3 | 90.20% | 75.40% |
| B31 N | 9.1 | 90.90% | 74.60% |
| B32 N | 9.2 | 92.00% | 75.90% |
| D11 N | 9.3 | 92.50% | 75.00% |
| D13 N | 9.3 | 92.40% | 73.60% |
| D22 N | 9.1 | 92.40% | 74.40% |
| D32 N | 9.4 | 91.60% | 74.20% |
| D34 N | 9 | 91.20% | 74.60% |

### RNAseq QC and Alignment

Initial quality control checks were performed by FastQC [13]. All samples showed mean quality scores > 30. Fastp v. 0.22.0 was used for adapter and end trimming and further quality control prior to alignment [14]. C57BL/6J and DBA/2J samples were aligned to release 108 of the Ensembl genome for C57BL/6J mice using STAR v 2.7.10b [15]. DBA/2J samples were also aligned to the DBA/2J genome provided by the European Nucleotide Archive (FASTA and GFF3, as below) using STAR v 2.7.10b. Aligned BAM files produced by STAR were further sorted by coordinate using Samtools v 1.6 [16]. Raw read counts for each BAM file were assigned and quantified using FeatureCounts (Samtools).

### D2 Annotation File Preparation

FASTA (GCA_921998315.2 2.fasta) and GFF3 (DBA_2J_v3.2.gff3) annotation files for the DBA/2J genome were downloaded (1/23/2023) courtesy of Thomas Keane at the European Genome-phenome Archive (European Bioinformatics Institute). Another GTF/GFF Analysis Toolkit (AGAT) was used to convert the GFF3 annotation file to a GTF file to be used for downstream sorting and DEXSeq count generation [17]. The converted GTF file was further modified by removing the “Parent” attributes that conflict with the “gene_ID” and “transcript_ID” attributes. GTF files do not contain parent relationships for the genes and transcripts contained within that are present in the GFF3 format. Removing the “Parent” attributes leaves the GTF file with the same attributes as a normal GTF file.

### Differential Gene Expression

Count files were analyzed for differential gene expression between B6 and D2 animals using the R package DESeq2 v 1.36.0 [18]. Low expressed genes that had median counts across all samples ≤ 0 were eliminated prior to analysis. Principal Component Analysis of the variance on the top 10,000 genes was run to identify significant sample outliers. Counts from B6 samples were compared to counts from D2 samples aligned with both the GRCmm39 reference genome and DBA/2J genome. Genes with a false discovery rate (FDR) < 0.05 were considered significantly altered and used in downstream analyses. Gene lists from each analysis were compared for overlap.

### DEXSeq Count Generation

A GFF annotation file containing collapsed exon counting bins was prepared from the UCSC GRCm39/mm39 GTF file using DEXSeq (v.1.42.10) with the python script *dexseq_prepare_annotation.py* [19]. The number of reads overlapping each exon bin was then counted using the DEXSeq Python script dexseq_count.py, the GFF file, and each sample’s BAM file. Differential exon usage (DEU) analysis was then carried out for the same contrasts studied in our DGE analysis using the DEXSeq R package standard analysis workflow. For D2 samples aligned to the D2 genome, the modified GTF file used (DBA_2J_v3.2_3_14_23_filtered.gtf) was prepared using the *dexseq_prepare_annotation.py* to generate the flattened GFF file used for count generation. The number of reads overlapping each exon bin was then counted using the DEXSeq python script *dexseq_count.py*, the flattened GFF files for both the GRCmm39 and DBA/2J genomes, and each sample’s BAM file. Differential exon usage (DEU) analysis was then carried out for the same contrasts studied in our DGE analysis using the DEXSeq R package standard analysis workflow.

### Gene Ontology and Semantic Similarity Analysis

Functional enrichment analyses for significant DGE and differential exon utilization (DEU) results were performed using ToppFun, available as part of the ToppGene suite of web-based applications [20]. Mouse gene symbols were submitted and analyzed for over-representation of genes that belong to Gene Ontology categories (molecular function, biological processes, and cellular component). Gene lists derived from the DESeq or DEXSeq analyses were filtered by FDR < 0.1 for Gene Ontology studies. Only categories with p-values less than 0.05 and possessing between 3 and 1000 total genes were considered. The webtool REVIGO was used for data reduction by semantic similarity using default parameters, and visualization of GO terms lists resulting from this analysis [21].

## Results

### Alignment to D2 Genome Improves Alignment Percentage

Here we conducted molecular analyses on strain-specific gene expression differences that occur as a result of differences in RNA-seq alignment. D2 samples were aligned to both the reference (B6) genome and a D2 genome and compared in separate analyses against B6 samples aligned to the reference genome. Aligning D2 samples to the D2 genome resulted in a significantly (p = 0.00272) higher percentage of reads aligned (93.82%) compared to the B6 genome (92.02%). However, the raw number of reads averaged across samples in each alignment was not statistically significant (**Table 2**, 133.66 M for B6 and 136.22 M for D2; p = 0.5354). Following alignment, counts were generated using *FeatureCounts* for downstream differential gene expression analysis. There was a significant decrease in the percentage of reads uniquely mapped in the count generation of D2 samples aligned to the D2 genome (72.3%) compared to the B6 genome count assignment (74.63%) (Table 1, p < 0.0001). Following count generation, two DESeq2 analyses were run comparing B6 samples to either the B6-aligned D2 samples or the D2-aligned D2 samples. Principal component analysis (PCA) of the top 10,000 most abundant genes by counts for B6-aligned D2 sample comparison showed that strains clustered together by 80% variance (**Figure 1A**). Alignment of D2-samples to the D2 genome improved this clustering to 98% variance (**Figure 1B**). These results establish that, despite a reduction in read assignment, the alignment of D2 samples to their parent genome results in greater specificity during sample alignment.

**Figure 1.**
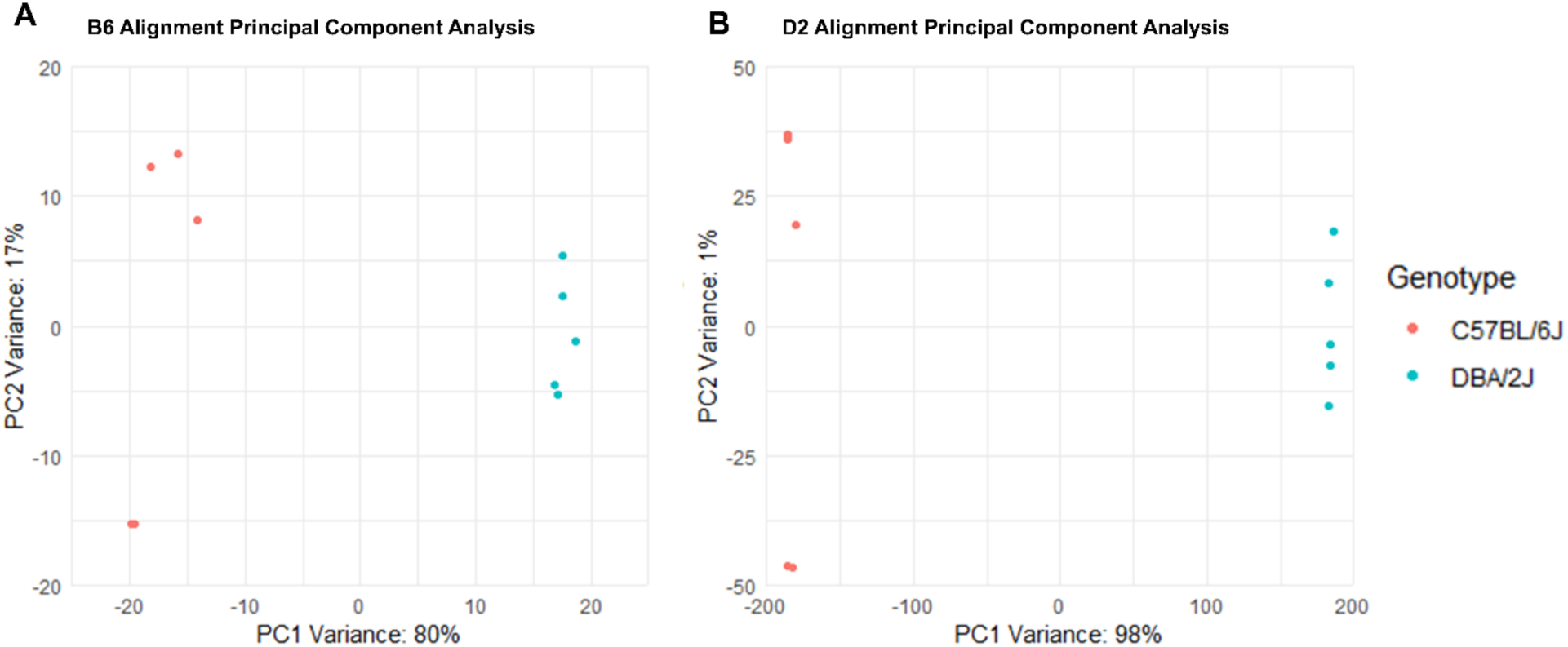
Principal Component Analysis (PCA) between Strain Alignments. The Top 10,000 genes identified by differential gene expression analysis (DESeq) was run to identify significant sample outliers.

**Table 2.** Comparison of C57BL/6J and DBA/2J Alignments and Count Generation. Alignment of D2 samples to their parent genome resulted in a significant increase in read alignment (p = 0.00272) but not read assignment (p = 0.5354). Read assignment showed a trend for a decrease in overall assignment. Data analyzed by Student’s T-Test.

|  | C57BL/6J Alignment |  |  |  | DBA/2J Alignment |  |  |  |
| --- | --- | --- | --- | --- | --- | --- | --- | --- |
| Sample | Reads Aligned (M) | % Aligned | Reads Assigned (M) | % Assigned | Reads Aligned (M) | % Aligned | Reads Assigned (Million) | % Assigned |
| D11 | 138.7 | 92.50% | 116.8 | 75.00% | 139.1 | 93.60% | 111 | 72.80% |
| D13 | 131.6 | 92.40% | 108.7 | 73.60% | 127.1 | 92.80% | 103 | 71.60% |
| D22 | 136.9 | 92.40% | 114.3 | 74.40% | 139.4 | 94.10% | 108.4 | 72.40% |
| D32 | 136.2 | 91.60% | 113.2 | 74.20% | 133.7 | 93.90% | 107.6 | 72.10% |
| D34 | 124.9 | 91.20% | 104.4 | 74.60% | 141.8 | 94.70% | 99.2 | 72.60% |
| AVG | 133.66 | 92.02% | 111.48 | 74.36% | 136.22 | 93.82% | 105.84 | 72.3% |

### D2 Alignment Increases the Number of Significant DEGs

Based on the observed increase in the variance between the B6 and D2 strains, as indicated by the PC1 of the principal component analysis, we performed two separate differential gene expression analyses using DESeq2. The first analysis compared DEGs between the B6 and D2 strains when both aligned to the reference genome. The second analysis compared DEGs between B6 and D2 strains when each strain was aligned to their strain-specific genome. A p-value cutoff of less than 0.05 was implemented for these analyses, with no filtering applied for log2 fold-change (LFC). In the first analysis with the reference genome alignment, a total of 6,210 genes exhibited differential expression between the two strains. Of these, 3,257 genes demonstrated a positive LFC, reflective of higher expression in the D2 strain while 2,953 genes showed a negative LFC, representative of higher expression in the B6 strain. In the second analysis where D2 mice were aligned to the D2 genome, differential gene expression analysis revealed that 10,777 genes were differentially expressed between the B6 and D2 strains. Among these genes, 5,463 displayed a positive LFC (higher expression in D2) while 5,314 had a negative LFC (higher expression in B6). The use of strain-specific alignment resulted in 4,567 more differentially expressed genes between the two strains.

### Functional Enrichment Analysis of Differentially Expressed Genes

While we showed the strain specific alignment improved specificity in overall read assignment and increased the number of DEGs identified between B6 and D2 mice, we wanted to examine the biological relevance of the genes being identified. The entirety of the 6,210 genes identified in the B6-alignment analysis and the 10,777 genes from the D2-alignment analysis were utilized as input for separate gene ontology over-representation analyses via ToppFun (**Figure 2**). The top 10 molecular function, biological processes, and cellular component categories were selected to represent our results. For molecular function, the B6-alignment analysis primarily featured categories related to cytoskeletal binding and actin binding. In contrast, the D2-alignment analysis emphasized categories such as zinc ion binding and SNARE binding (**Figure 2A**). For biological processes, the B6-alignment analysis highlighted significant categories such as gliogenesis, glial cell differentiation, and glial cell development. Conversely, the D2-alignment analysis exhibited a higher representation of genes associated with translation at various synaptic locations: presynapse, postsynapse, and synapse (**Figure 2B**). Cellular components enriched for the B6-alignment analysis included the synapse and golgi apparatus, while the D2-alignment analysis emphasized categories for the cytosolic ribosome and vesicle membranes (**Figure 2C**).

**Figure 2.**
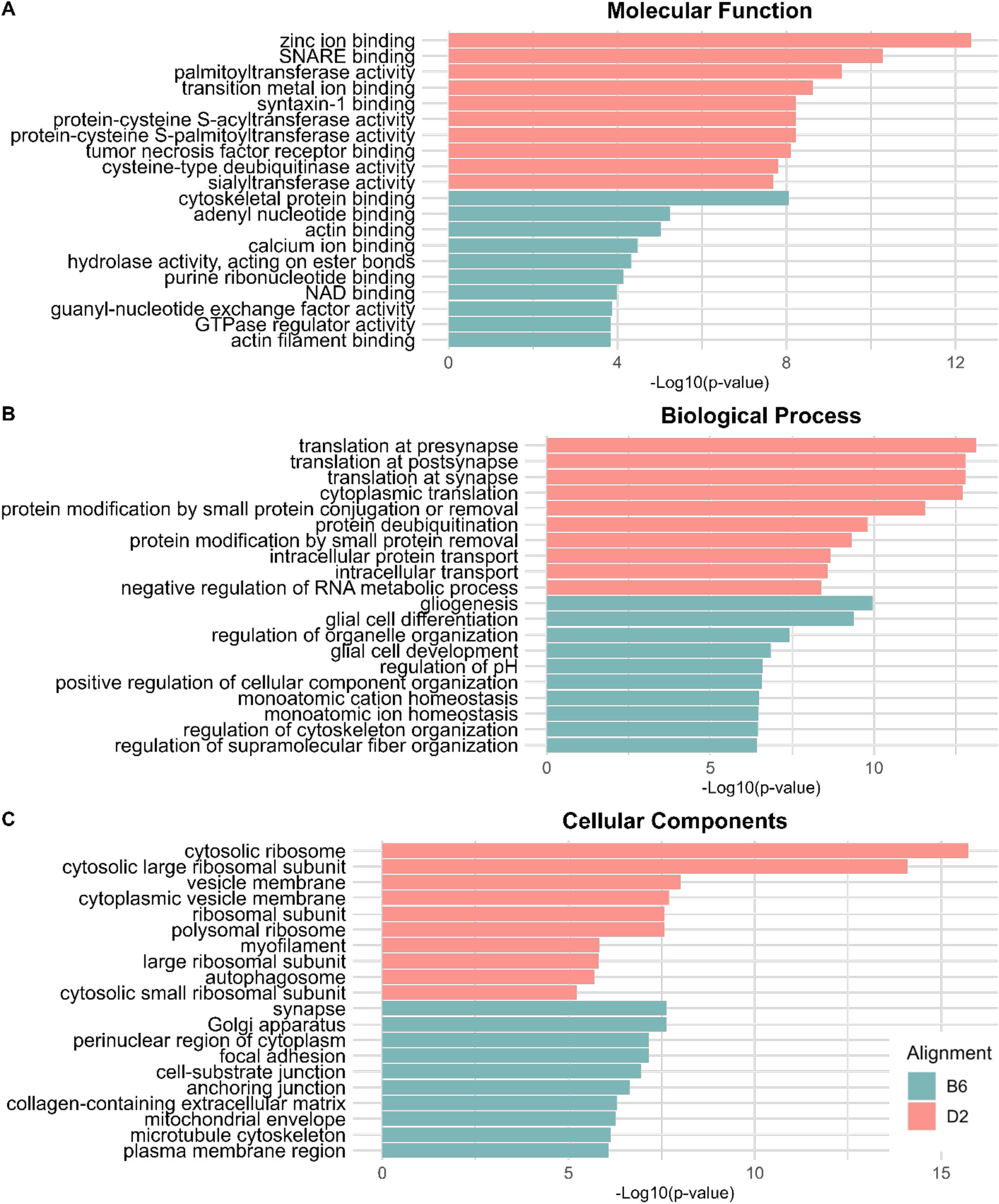
Differential Gene Expression Gene Ontology. Differentially expressed genes were compared between B6 mice and D2 mice aligned either to the B6 or D2 genome and used for functional enrichment analyses. Significance is Log-base-10 of the p-value.

### Alignment Exclusive Differential Gene Expression

After comparing overall differences between the two analyses, we further investigated the characteristics of overlapping differential expressed genes as well as identifying categories of genes unique to the two alignment analyses (**Figure 3**). All genes which exhibited a sign change for LFC were excluded from the following analysis. 620 DEGs were significant only in the B6 alignment while 5,206 DEGs were significant only in the D2 alignment and 5571 genes overlapped between the two alignments (**Figure 3B**). Gene Ontology analysis was used for functional surveys of the DEGs. The DEGs unique to the B6 alignment only generated cellular component categories. Genes identified only in the B6 alignment had functional clusters related to proton-transporting two-sector ATPase complex and eukaryotic translation initiation factor 3 complex. The D2-alignment exclusive DEGs top (**Figure 3C**) Molecular Function categories included zinc ion binding, deubiquitinase activity, and transcription regulatory activity. Top Cellular Component categories included the cytosolic ribosome and cytosolic large ribosomal subunit. Top Biological Processes included modification-dependent protein catabolic processes and ubiquitin-dependent protein catabolic processes.

**Figure 3.**
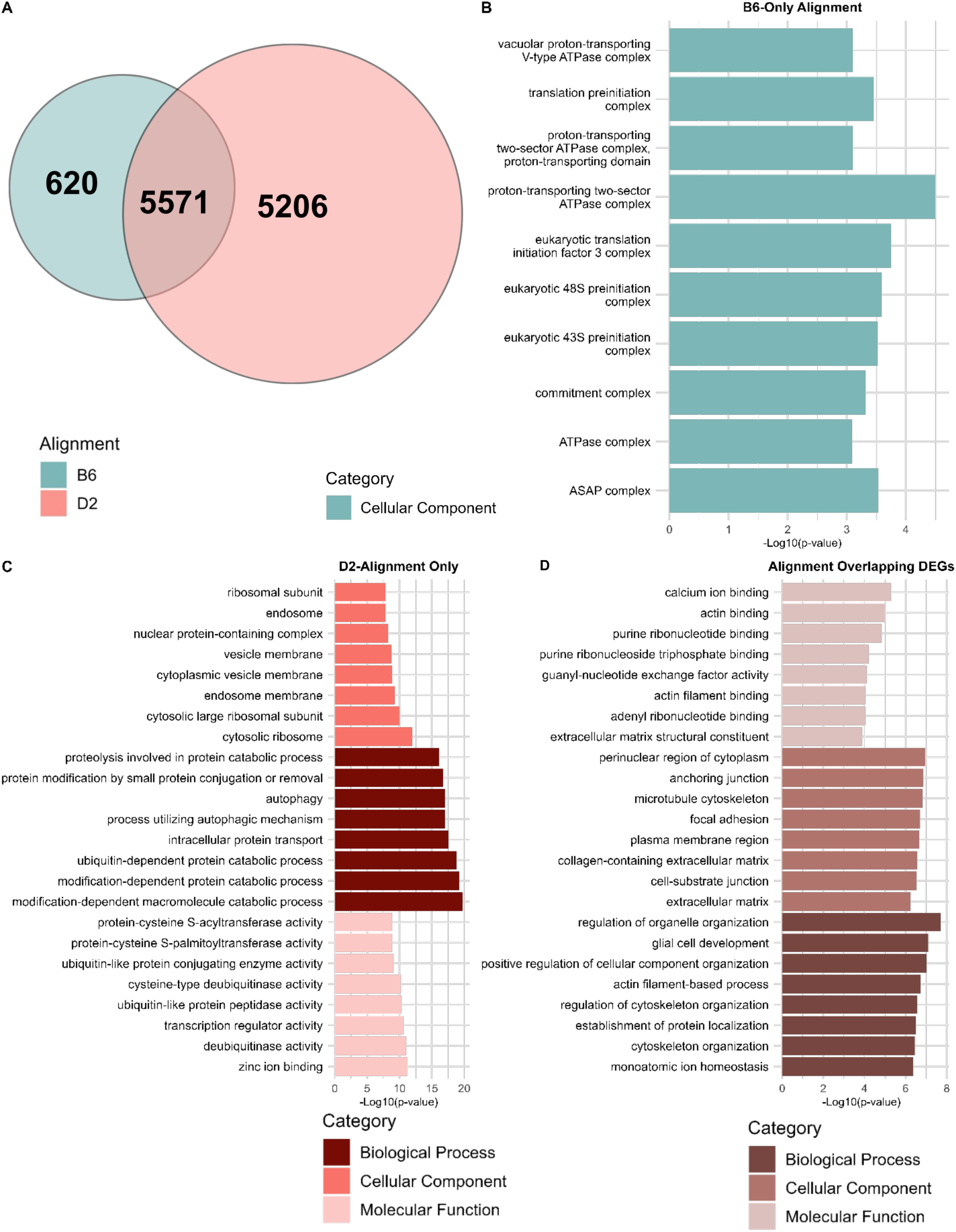
Gene Ontology of overlapping DEGs and Independent Alignment DEGs. **(A)** Venn Diagram of number of DEGs identified between B6 and D2 mice when the D2 mice are aligned to the B6 genome (green) or the D2 genome (pink). **(B)** Gene ontology of B6-aligned DEGs (n = 620). **(C)** Gene Ontology of D2-aligned DEGs (n = 5206). **(D)** Gene Ontology of overlapping DEGs identified by both analyses (n = 5571). All gene ontology results are presented as the negative Log-based-10 of the p-value.

### Overlapping DEGs

Functional enrichment analysis of the 5,571 DEGs identified as overlapping between the B6 and D2 alignment analyses (**Figure 3D**) largely showed top categories unique from those seen with the individual alignments. Over-representation analysis identified Molecular Functions such as cytoskeletal protein binding, nucleotide binding; Cellular Components such as structural proteins and the extracellular matrix; and Biological Processes such as glial cell differentiation and development and cytoskeleton organization. The cytoskeleton, which is comprised of actin filaments, microtubules, and intermediate filaments, is a key player in maintaining cellular structure and organization [22]. The consistent identification of these genes in both analyses suggests that the genes associated with the cytoskeleton exhibit significant conservation across strains, with minimal polymorphisms affecting the expression patterns revealed by sequencing.

### D2 Alignment Alters Directionality of LFC

In the comparison of these two alignments, a total of 479 genes were observed to exhibit a change in sign for their Log Fold Change (LFC) values indicating a shift in which strain exhibits higher expression of the gene. Among these, 290 genes exhibited a sign change in LFC values from a positive to a negative LFC, suggesting these genes now have higher expression in B6 samples. Conversely, 189 genes demonstrated a shift from negative to positive LFC, indicative of now higher expression in the D2 samples (**Figure 4**, **Table 4**). Subsequent Gene Ontology (GO) analysis was performed on both gene sets. However, significant GO categories were only generated from the set of genes showing a change to upregulated expression in B6 samples. The top categories under Molecular Functions were identified as structural constituents of the ribosome and inhibitors of ubiquitin ligase or protein transferase activity. This aligns with overrepresented Cellular Components which include the cytosolic ribosome and ribonucleotide protein complex. The predominant Biological Processes involved genes associated with cytoplasmic translation, ribosomal biogenesis and organelle assembly (**Figure 4B**). In contrast to DEGs that overlapped between the two analyses, ribosomal protein DEGs whose expression changed to being higher in the B6 animals highlight an area where misalignment may be frequently missed.

**Figure 4.**
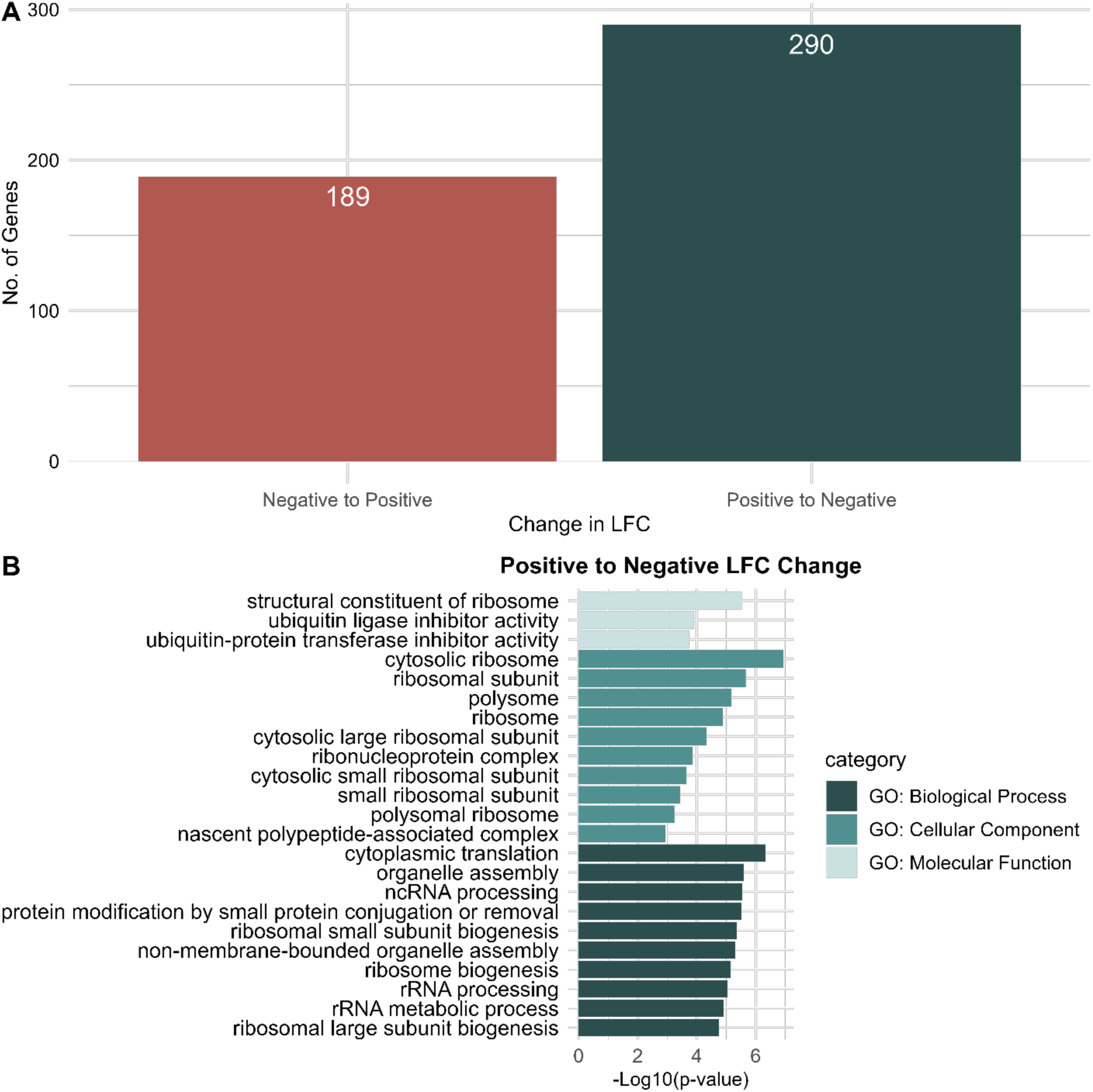
Gene Ontology of DEGs with Different LFCs between Alignment. **(A)** Representative bar graph of the number of DEGs with significant shifts in log fold change (LFC). **(B)** Gene Ontology analysis of DEGs where the LFC changed from positive (higher D2 expression to higher B6 expression).

### D2 Alignment Increases Differential Exon Utilization

We performed two differential exon utilization analyses in order to determine the effect of strain specific alignment on exon level expression. The first analysis compared differentially utilized exons between the B6 and D2 strains with both aligned to the reference genome. The second analysis compared differentially utilized exons (DUEs) between B6 and D2 strains when each strain was aligned to their strain-specific genome. The B6 aligned analysis identified 21,223 differentially utilized exons across 6,650 genes, while the D2 aligned analysis identified 81,206 exons across 13,521 genes (**Figure 5A**). 14,245 differentially utilized exons across 5,828 of those genes overlapped between the two analyses. In order to further understand the functions of genes with differentially utilized exons, the 6,650 genes identified in the B6 aligned analysis and the 13,521 genes identified in the D2 aligned analysis were separately used as input for functional over-representation analysis. REVIGO semantic similarity analysis was used to cluster gene ontology terms within the biological processes, molecular function, and cellular component categories in order to reduce the complexity of comparisons (**Figure 5B-D**). The functional enrichment categories found in the B6 aligned analysis were strongly conserved in the D2 aligned analysis, with 85.20% for biological processes, 61.19% for molecular functions, and 85.26% for cellular components. However, despite the increased number of DUEs with the D2 alignment, the p-values for over-represented REVIGO groupings were consistently lower than with the B6 alignment.

**Figure 5.**
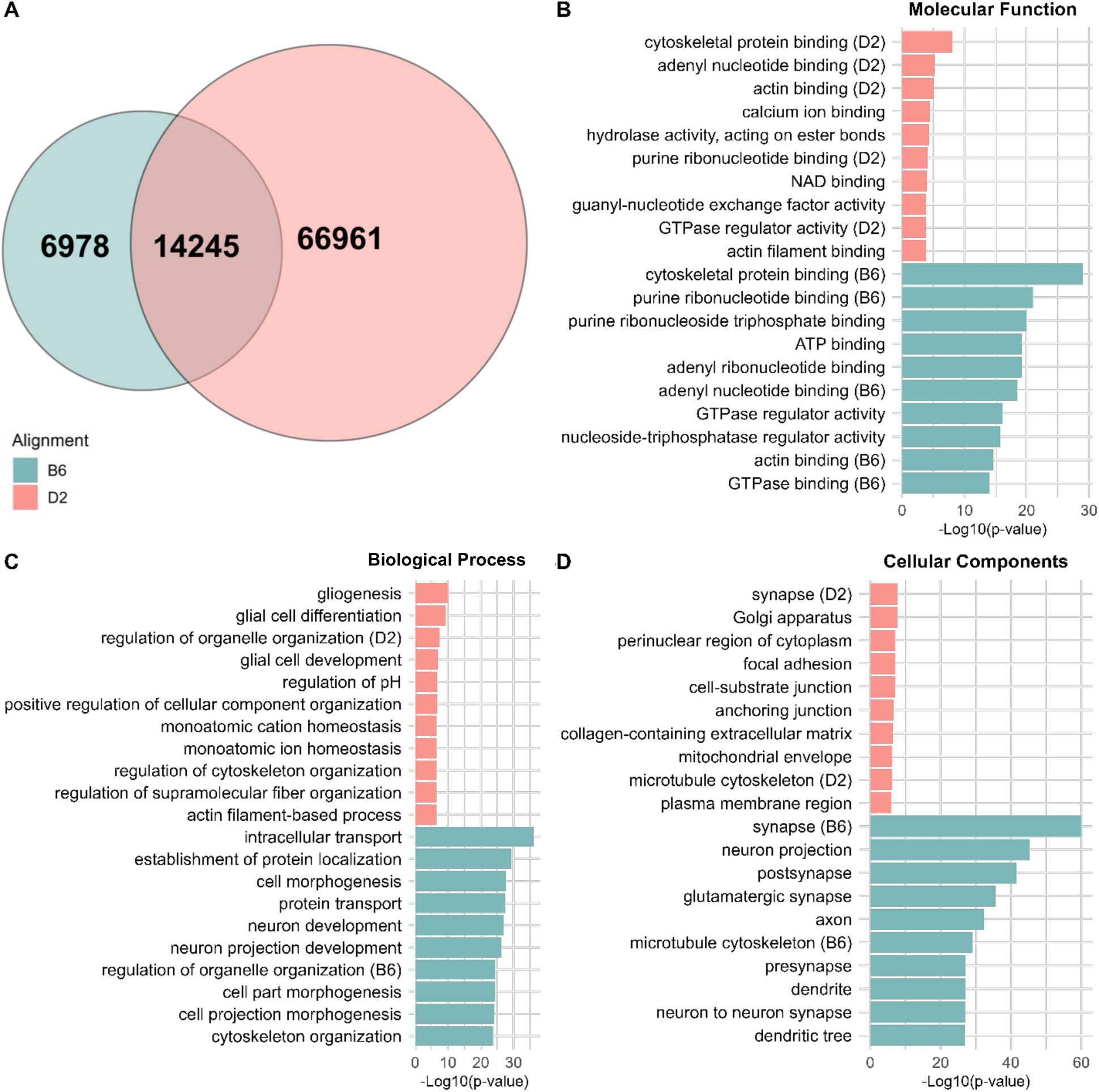
DEGs and Gene Ontology of DEXSeq Results. **(A)** Venn Diagram representing all differentially utilized exons identified using the B6 and D2 alignments. Top 10 gene ontology categories for each alignment are divided by **(B)** Molecular Function, **(C)** Biological Process, and **(D)** Cellular components. Significance is represented by Log-based-10 of the p-values.

The largest semantic groupings were conserved in each, with the top 4 semantic groupings for B6 aligned biological processes being regulation of organelle organization, intracellular transport, mRNA processing, and cytoskeleton organization. The top 4 in the D2 aligned biological processes being neuron projection development, organonitrogen compound catabolic process, regulation of organelle organization, and intracellular transport. The top 4 semantic groups for B6 aligned molecular functions were cytoskeletal protein binding, single stranded DNA binding, phosphotransferase activity – alcohol group as acceptor, and calcium ion transmembrane transporter activity. In D2 aligned molecular functions the top 4 were cytoskeletal protein binding, phosphotransferase activity – alcohol group as acceptor, zinc ion binding, and ribonucleoside triphosphate phosphatase activity. The B6 aligned semantic similarity analysis identified 3 semantically similar clusters; every other term was distinct enough for its own individual cluster. Those top 3 groups were microtubule cytoskeleton, synapse, and neuron projection, while the top 4 in the D2 aligned analysis were microtubule cytoskeleton, synapse, neuron projection, and plasma membrane region. Many of the individually clustered terms in the B6 cellular components were conserved in the D2 aligned analysis.

## Discussion

The process of alignment in RNA-seq is a critical step in accurately representing gene expression levels that reflect biological reality. Misalignments can lead to the inaccurate quantification of gene expression, false discovery of novel transcripts, and erroneous interpretation of alternative splicing events. To our knowledge, the studies here provide the first genomic analysis examining the role of RNA-seq alignment on differential gene expression and subsequent functional enrichment analyses. Using the most extensive D2 genome available, we demonstrated that alignment of D2 samples to their parent genome resulted in increased read alignment percentage and subsequently, an increased number of differentially expressed genes and exons.

Our studies showed that alignment of DBA/2J samples to their parent genome improved alignment percentages by on average 1.8%. This is consistent with previous results from Gobet et. al 2022 showing that alignment of D2 samples to the D2 assembly improved the percentage of uniquely mapped reads by 0.03-3.4% [4]. For both assembly alignments, our alignment percentages are consistent with the alignment rates published by the makers of the STAR aligner [15]. Despite improvements in read alignment, the D2 alignment produced a lower percentage of assigned reads than the B6 aligned during the count generation step. This could be attributed to several factors, including the complexity of the genomic regions or genetic variations between the samples and the D2 reference genome. However, the most likely explanation is that the D2 reference genome sequence is not as completely aligned compared to the B6 reference genome and is less well annotated. Regions that aren’t well represented in the D2 reference genome would cause their associated reads to be assigned at a lower rate or not at all. However, the percentage of read assignment is still high, being within 2% of the B6 alignment. Despite these decreases in overall read assignment, principal component analysis of the normalized counts demonstrated an improvement in PC1 indicating that 98% of the variance in the expression of the Top 10,000 genes is explained by strain differences. The benefits of aligning to the D2 genome, such as increasing the future analyses’ ability to detect SNPs and small indels, and potential allele specific expression differences, outweigh the slight decrease in assignment percentage moving forward.

It is evident that strain-specific alignment is important for increasing the specificity of read assignment. There are currently few studies that extend the characterization of alignment to accurately understanding the biological processes that are represented by differentially expressed genes. Our functional enrichment analysis underscores the observation that the detection of differentially expressed structural protein genes remains highly conserved, irrespective of the alignment annotation employed. This may be reflective of either a lower density of non-synonymous polymorphisms or the superior quality of annotation for these genes given their significance in cell structure and function like the cytoskeletal proteins. Unlike the conserved detection of structural protein genes, the DEGs unique to the B6 and D2 reference genomes show a divergent representation of functional categories. DEGs only found when aligning to the B6 reference genome were over-represented for genes involved in translation complexes and the ATPase complex. Conversely, DEGs identified exclusively by the D2 genome alignment were over-represented for genes encoding ribosomal proteins and proteins involved in ubiquitin-dependent processes. Though 5 million SNPs have been identified to differ between B6 and D2 mice, further analysis is needed to further examine the distribution of these SNPs based on biological function and relevance.

It is clear from our analyses that the directionality change of a gene’s LFC suggests that the aligning of the D2 strain to the reference genome may result in the misalignment of genes, as evident in the case of ribosomal protein genes identified in this analysis. It may be that, in contrast to the overlapping genes, genes with a higher density of polymorphisms may ultimately result in the alignment of reads to the wrong gene or not aligning at all. Highly structurally conserved gene groups such as ribosomal proteins might be particularly susceptible to this issue. However, current analysis software programs are not equipped to make comparisons between groups that use different reference genomes. This may result in falling to characterize novel splicing events that are specific to the D2 strain.

In an attempt to characterize splicing events between the alignments, we performed a differential exon utilization analysis. In this alignment comparison, we saw an increase in both the number of differentially utilized exons and the genes that contained those exons. The D2 aligned analysis added novel differentially utilized exons while conserving 87.64% of the exons identified by the B6 aligned analysis. However, this could be due to factors outside of the strain specific alignment. The overall quality of an annotation is crucial in accurately defining exonic and intronic boundaries that sequencing reads align to. A lower quality annotation may inaccurately define the chromosomal boundaries of exons and introns, and these annotations tend to lack information regarding alternative splicing junctions and events. A higher quality annotation includes more detailed characterizations of alternative splicing events and transcript variant identification that are important in characterizing expression for an exon-level analysis. This greater degree of information and specificity reduces the number of false positive read assignments and annotation. Given the B6 genome annotation is the reference mouse genome, it has the highest quality annotation available for mouse genomics and is frequently updated. While our D2 annotation more accurately reflects the D2 genome, it may lack the depth of information the B6 genome provides. We saw a high level of conservation in the functional enrichment categories identified by the D2 aligned analysis, with 96.91% of functional enrichment categories identified by the B6 aligned analysis also being identified by the D2 aligned analysis.

It has been repeatedly argued that the exclusive use of B6 mice or those on a B6 background in the investigation of gene expression and function may result in misidentifying biologically relevant pathways [23; 24]. This exclusivity extends into the use of the B6 genome as the reference genome, which is highly used in the alignment of other strains, including the DBA/2J strain. There is a clear need for the rapid improvement of strain-specific reference genomes with greater annotation depth comparable to that of the current reference genome. To our knowledge, current analysis software for the investigation of alternative splicing is unable to perform such analyses when samples are aligned to different references given their dependence on BAM inputs. Therefore, we are hindered in our abilities to accurately characterize gene expression regulation and transcript composition differences. The advent of full characterizing the genomes other commonly used inbred strains to the extent of the C57BL/6J reference assembly will also generate the need for new downstream analysis software to perform such analyses.

## Acknowledgements

The authors thank the members of the Miles laboratory for their comments and insight during the completion of these studies. The data included in this study was generated at the Genomics Core facility at Virginia Commonwealth University. This work was supported by grants F31AA030727 (ERG) and P50AA022537 (MFM) from the National Institute on Alcohol Abuse and Alcoholism (NIAAA).

## Notes

https://www.ncbi.nlm.nih.gov/gds/?term=GSE274854

### Competing Interest Statement

The authors have declared no competing interest.

